# Using CCDC6 immunostaining in conjunction with the RAD51 HRD assay as a novel approach to expand PARPi treatment eligibility in HGSOC patients

**DOI:** 10.1101/2025.07.29.667411

**Authors:** Daniela Criscuolo, Francesco Merolla, Benedetta Pellegrino, Luca Russolillo, Ilaria De Benedictis, Daniela Califano, Rosaria Catalano, Carmela Baviello, Silvia Varricchio, Sabrina C Cecere, Camilla Nero, Eleonora Palluzzi, Dyonissios Katsaros, Ettore D Capoluongo, Giovanni L Scaglione, Sergio Marchini, Daniela Russo, Anna Spina, Laura Arenare, Francesco Morra, Maria Marotta, Marialuisa A Vecchione, Alexandra Ingallinella, Francesco Perrone, Sandro Pignata, Angela Celetti

**Author notes:** Last & Corresponding Author: Angela Celetti, MD, PhD, CNR – Institute of Endotypes in Oncology, Metabolism and Immunology “G Salvatore” -IEOMI- Naples, Italy. **Declaration of competing interests** The authors have no competing interests to declare.

## Abstract

**Purpose:** HGSOC patients with BRCA1/2 mutations show HRD and PARPi sensitivity. Notably, HRD and PARPi response can occur without BRCA mutations, suggesting other factors are involved. Loss of CCDC6 function can lead to HRD and PARPi sensitivity in HGSOC cells, making CCDC6 a potential therapeutic target and biomarker. Three CCDC6 missense mutations in HGSOC prompted investigation into their impact on HRD. Analyzing CCDC6 expression, localization and HRD data in the MITO16A trial aims to clarify the CCDC6-HRD relationship in a large cohort.

**Experimental Design:** The biochemical and morphological effects of CCDC6 mutants on the native protein were examined using pull-down assays and immunofluorescence. HR-reporter and cell viability assays determined the impact of these mutants on HRD and PARPi sensitivity. CCDC6 histochemical score and intracellular-localization were assessed in MITO16A samples after immunostaining and digitalization.

**Results:** CCDC6-mutated isoforms act as dominant-negative, preventing native CCDC6 nuclear translocation, disrupting RAD51 foci and HR-repair, and increasing PARPi sensitivity. In the MITO16A patient sample set, 66 of 185 (35%) showed barely detectable CCDC6 or nuclear exclusion (“CCDC6-inactive”). CCDC6 impairment in these “CCDC6-inactive” samples was associated with HRD in 75% (30/40) of suitable samples analyzed by the RAD51 test and in 52% (34/65) of suitable samples analyzed by genomic HRD testing, even in the presence of wild-type BRCA1/BRCA2 genes.

**Conclusion:** The association between CCDC6 inactivity and HRD, both at genomic and functional level, occurred even in presence of wild-type BRCA1/BRCA2 genes, suggesting that CCDC6 may play a crucial role in DNA repair pathways independent of these well-known genes.

## Introduction

The most common and aggressive histotype of ovarian cancer is represented by the high-grade serous ovarian carcinoma (HGSOC), one of the main causes of death from gynecological malignancies.^1^ Tumors carrying BRCA1/BRCA2 mutations exhibit a phenotype known as BRCAness, characterized by deficiency in DNA Double Strand Breaks (DSBs) Repair by Homologous Recombination (HR).^2^ This phenotype can be therapeutically targeted through a synthetic lethal strategy, which involves the use of poly-(ADP-ribose)-polymerase inhibitors (PARPi).^3, 4^ Notably, PARPi also prolong progression-free survival (PFS) in 50-60% of BRCA1/BRCA2 wild-type HGSOC patients, due to other underlying HR repair deficiencies.^5-7^

In 2020, Lynparza gained FDA/EMA approval for maintenance treatment in HRD-positive ovarian cancer, highlighting HRD’s role in PARPi efficacy.^8, 9^ European guidelines now recommend HRD testing for all advanced OC patients, beyond BRCA status, since various HRD assays exist ^9-12^. RAD51 foci assessment in tumors is a promising method to predict PARPi response, reflecting HR efficiency. ^13-15^ Low RAD51 scores correlate with PARPi sensitivity in breast, prostate, and HGSOC cancers. ^16-18^ Further research into other HR-related molecular events and genes is needed to improve precision medicine and expand the range of patients who can benefit from PARPi therapy.

In various cancers, the CCDC6 (Coiled Coil Domain Containing 6) tumor suppressor protein often loses its function due to genetic alterations, including translocations, somatic mutations, and changes in protein levels.^19^ Following DNA damage, CCDC6 moves to the nucleus, where it contributes to HR repair by regulating H2AX phosphorylation through its interaction with the PP4c protein phosphatase.^20^ We have recently shown that loss of CCDC6 in HGSOC cells disrupts HR repair and induces sensitivity to PARPi, suggesting CCDC6 as a potential biomarker for PARPi response.^21^ Notably, three CCDC6 missense mutations (L217P, A226S, P442S) have been identified in approximately 3% of primary HGSOC tumors (https://cancer.sanger.ac.uk/cosmic).

Our study characterized how the mutated isoforms of the CCDC6 protein alter the location of the native CCDC6 protein within the cell and how this affects the protein’s role in HR repair. A key finding was that when CCDC6 is excluded from the nucleus, its function in HR repair is compromised. This impairment of HR repair due to CCDC6 nuclear exclusion leads to increased sensitivity to PARPi in HGSOC cells. Analysis of 185 HGSOC patient samples (FFPE tissues) from the MITO16A trial, which had already undergone genomic and functional HRD testing^21^, showed a correlation between loss of CCDC6 (either due to low expression or mislocalization) and HRD. This association with HRD was particularly strong when HRD was assessed using the RAD51 assay, suggesting this assay may be especially sensitive to CCDC6-related HR impairment.^22, 23^

Given the current limitations in accurately assessing HRD, combining CCDC6 immunostaining with both genomic and functional HRD tests offers a promising approach to improve prediction of PARPi treatment response in HGSOC. This combined strategy could be a valuable tool for identifying HRD-positive patients who are likely to benefit from PARPi therapy, and importantly, it may broaden our understanding of HRD beyond BRCA1 and BRCA2 mutations.

## Material and methods

### Cell lines, drugs and chemicals

All the cells used in the study were authenticated using STR profiling and were routinely checked for mycoplasma contamination using the PCR Mycoplasma Detection Kit (ABM), according to the manufacturer’s protocol. HeLa Kyoto cells (RRID: CVCL_1922), harboring the S-tag-GFP-CCDC6 construct, were generated by Ina Poser in the laboratory of Anthony Hyman and were grown in DMEM (high glucose), 1% FCS, 1% penicillin/streptomycin (Gibco, Paisley, UK). ^24, 25^ Ovarian cancer cell lines OVCAR3 (RRID: CVCL_0465) and OV90 (RRID: CVCL_3768) were purchased by ATCC and maintained in RPMI and DMEM, respectively, with 10% FBS, 1% L-glutamine, and 1% penicillin/streptomycin (Gibco, Paisley, UK). Drugs and chemical: Olaparib (AZD2281, S1060-SelleckChem), cisplatin (P4394-Merck Millipore), and Hoechst-33258 (94403-SIGMA-Aldrich).

### Plasmids and transfection

pcDNA4ToA/myc-his-CCDC6 plasmid (V1030-20, InVitrogen Corporation, San Diego, CA, USA) were transfected using FuGene HD (Promega). Myc-CCDC6 mutants (A226S, L217P, and P442S) were generated from the wild-type pcDNA4ToA/myc-his-CCDC6 template using the Quick-Change Site-Directed Mutagenesis Kit (Agilent). CCDC6 shRNA (pLKO.1 puro, RRID: Addgene_8453) was purchased from Sigma-Aldrich. The generation of the stable silenced cell lines (OVCAR3 and OV90) have been described elsewhere. ^21^ The pDRGFP reporter plasmid (RRID: Addgene_46085) and the pCAGGS-I-SceI (RRID: Addgene_26477) have been utilized for the Homologous recombination transient assay, as reported below.

### Reagents and antibodies

Antibodies against: CCDC6 (Abcam Cat# ab56353, RRID: AB_940832) and (Atlas Antibodies Cat# HPA019051, RRID: AB_1846164), Rad51 (Santa Cruz Biotechnology Cat# sc-8349, RRID: AB_2253533) and (Abcam Cat# ab133534, RRID: AB_2722613), Myc (Santa Cruz Biotechnology Cat# sc-40, RRID: AB_627268), GFP (Cat# 632376, BD Living Colors, Takara Bio), HA (Santa Cruz Biotechnology Cat# sc-805, RRID: AB_631618), γH2AX, (Merck Millipore Cat# 05-636, RRID: AB_309864), Tubulin (Sigma-Aldrich Cat# T6557, RRID: AB_477584) were utilized for biochemical analysis and immunohistochemistry. Bio-Rad provided anti-mouse and anti-rabbit secondary antibodies (Bio-Rad Cat# 170-5047, RRID: AB_11125753; Bio-Rad Cat# 170-5046, RRID: AB_11125757), for western blot analysis. Abcam provided fluorescent anti-mouse and anti-rabbit secondary antibodies (Abcam Cat# ab150116, RRID: AB_2650601, Abcam Cat# ab150113, RRID: AB_2576208, Abcam Cat# ab150077, RRID: AB_2630356).

### Protein extraction and Western Blot Analysis

Total protein extracts and western blotting procedures have been previously described. ^21^

### Immunofluorescence staining

Following genotoxic stress exposure [etoposide (10μM)] in transfected and control cells immunofluorescence has been performed, as reported. ^25^ Cells with ≥ 5 distinct γH2AX or RAD51 foci were considered positive. The percentage of positive nuclei was calculated from a minimum of 250 analyzed nuclei per sample.

### Homologous recombination transient assay

OVCAR3 and OV90 cells transfected with myc-tagged CCDC6 mutants (A226S, L217P, P442S), or empty vector (EV), were plated in 12-well plates and transfected with the pDR-GFP reporter alone (negative control) or with pCAGGS-I-SceI. After 48 hours, cells were collected and analyzed by FACS using a Miltenyi MACSQuant Analyzer 10 - Flow Cytometer (Miltenyi Biotec).

### Sensitivity Test and Design for Drug Combination

The CellTiter 96 AQueous One Solution Cell Proliferation Assay (Promega) was used to determine drug IC50 values, as described. ^21^ CompuSyn software was used to analyze drug combinations, calculating the combination index (CI): CI = 1 (additivity), CI > 1 (antagonism), and CI < 1 (synergism).

### IHC analysis

185 HGSOC tissue samples from the MITO16A trial (NCT01706120 - National Cancer Institute, Naples) were processed for IHC as described before.^23^ 5 μm sections of formalin-fixed, paraffin-embedded (FFPE) tissues were stained with H&E or anti-CCDC6 antibody (HPA-019051, Sigma-Aldrich). CCDC6 immunoreactivity (staining intensity, fraction of stained cells, subcellular localization) was visually annotated. Digital analysis of scanned IHC slides (Leica AT2, 40x) using QuPath quantified CCDC6 expression via a validated pipeline (watershed algorithm for cell detection, DAB intensity thresholds for classification into 0, 1+, 2+, 3+ tiers), to calculate an H-score to quantify CCDC6 expression levels^23^. The median H-score across all cases was 211, and the mean was 204. Subcellular localization was also annotated. QuPath (RRID:SCR_018257) pipeline details are available upon request.

Based on the H-score and the subcellular localization of the protein, cases were categorized into Inactive (defined by either an H-score < 173 or complete nuclear exclusion of CCDC6) or Functional (H-score > 173 with diffuse or predominantly nuclear staining).

Figure S1A presents sample size (N), range (min-max), average, and median H-scores for functional and inactive categories. Quartile distribution is shown in Figure S1B.

The immunohistochemical (IHC) analysis of CCDC6 expression (H-score > 173) and localization (diffuse or mainly nuclear) of the “CCDC6 functional” MITO16A patient set (N = 117) is shown in the representative images of Figure S2.

### Statistical analysis

Statistical analysis was performed using GraphPad Prism v10 software (RRID: SCR_002798). All data are presented as Mean ± SEM. Unpaired Student’s *t* test was used for comparative analysis between two data groups. The following asterisk rating system for *P* value was used: ^*^*P* < 0.05; ^**^*P* < 0.01; ^***^*P* < 0.001; ^****^*P* < 0.0001.

## Results

### CCDC6 mutated isoforms interact with CCDC6 wild-type protein inducing its nuclear exclusion

CCDC6, a gene frequently mutated in cancer, shows 877 alterations across tumors, with missense substitutions being most common (https://cancer.sanger.ac.uk/cosmic). Around 3% of ovarian carcinomas (OC) in COSMIC have CCDC6 mutations. We focused on three missense mutations (L217P, A226S, P442S) in HGSOC patients, which, despite their location, could affect CCDC6 function and localization, similar to what’s seen in NSCLC.^25^

To check if CCDC6 mutants interact with the normal CCDC6 protein, we introduced myc-tagged mutant versions into HeLa cells expressing GFP-S-tag-CCDC6 (wild-type) and control HeLa cells. Using S-protein resin to pull down the S-tagged wild-type CCDC6, we found that all tested myc-tagged CCDC6 mutants (L217P, A226S, P442S) could interact with the native protein (Figure 1A, B). Confocal microscopy revealed that while the GFP-CCDC6 wild-type protein showed both nuclear and cytosolic localization (Figure 1C, c), the myc-tagged mutants were mainly cytosolic (Figure 1C, g,k,o). The merged images further confirmed these observations (Figure 1C, d,h,l,p)

**Figure 1.**
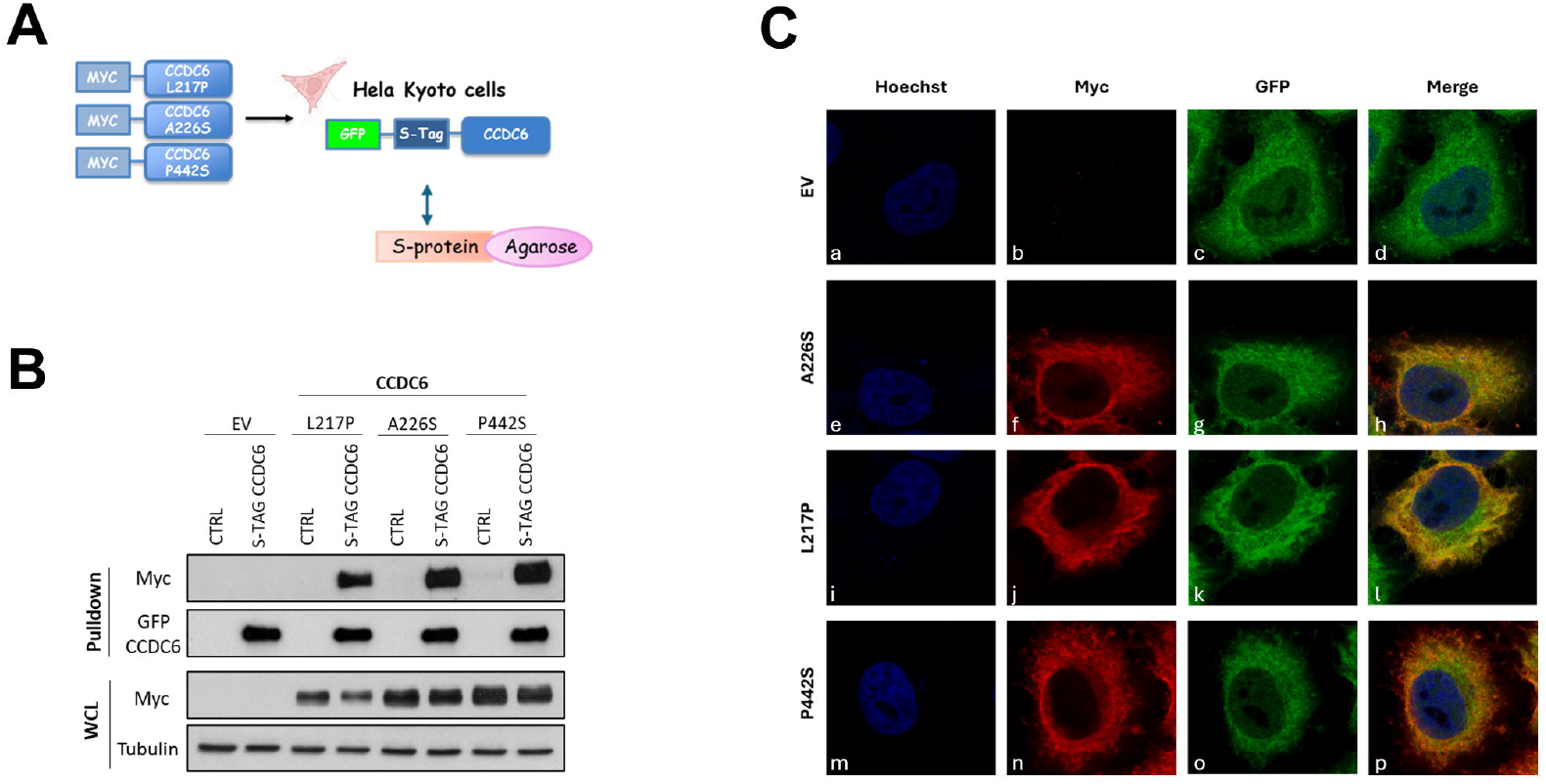
CCDC6 mutated isoforms form heterodimers with the CCDC6 wild-type protein and affects its intracellular distribution. (A) Schematic representation of the GFP-S-tag-CCDC6 construct and the S-protein agarose resin. (B) S-tag pull down of HeLa-Kyoto GFP-S-tag-CCDC6, and of HeLa-Kyoto wild type, transfected with the myc-tagged CCDC6 mutants L217P, A226S and P442S expression vectors, or with the empty vector (EV). Isolated proteins were immunoblotted with the anti-myc and anti-GFP antibodies. The immunoblots of the whole cell lysates (WCL) with anti-myc, for transfection control, and anti-tubulin, as loading control, are shown at the bottom of the panel. (C) Immunofluorescence images of HeLa Kyoto GFP-S-tag-CCDC6 transfected with the myc-CCDC6 mutated isoforms L217P, A226S and P442 expression vectors (e-p), or the myc empty vector (EV) as control (a-d). Nuclei are stained with the non-intercalating dye Hoechst (Blue-channel). CCDC6 wild type is shown as the endogenous GFP-S-tag-CCDC6 (Green-channel). The mutated isoforms were visualized by the anti-myc antibody (Red-channel).

This indicates that the CCDC6 mutants can heterodimerize with the wild-type protein and prevent its nuclear translocation. Thus, these CCDC6 mutants exhibit a dominant-negative effect on wild-type CCDC6 localization, causing it to be primarily cytosolic.

### CCDC6 mutated isoforms reduce γH2AX and RAD51 foci formation impairing DNA repair by homologous recombination pathway

When DNA damage occurs, CCDC6 is phosphorylated by ATM, moves to the nucleus, and aids HR repair by regulating H2AX phosphorylation through its interaction with PP4c. ^20, 26^ Since CCDC6 depletion impairs γH2AX foci formation and HR-mediated DSBs repair in HGSOC cells, ^27, 21^ we investigated the impact of CCDC6 mutants on the DNA damage response in BRCA1/2 wild-type HGSOC cell lines (OVCAR3 and OV90). Immunofluorescence analysis showed that overexpressing CCDC6 mutants significantly reduced γH2AX foci formation after treatment with genotoxic stress (Figure S3 A-B). This suggests that the mutants’ inability to translocate to the nucleus disrupts DNA damage signaling.

RAD51 is essential for accurate HR repair at DNA breaks. After damage, RAD51 forms nuclear foci at these sites, indicating active HR. ^28^ To understand CCDC6’s role in HR, we examined RAD51 foci formation in BRCA1/2 wild-type HGSOC cells with either CCDC6 mutant overexpression or stable CCDC6 depletion. Both CCDC6 loss and mutant overexpression led to reduced RAD51 recruitment to nuclear foci compared to controls (Figure 2A-D). Furthermore, both conditions showed decreased RAD51 protein levels in western blot analysis (Figure S3 C-D), suggesting CCDC6 is likely involved in the HR pathway.

**Figure 2.**
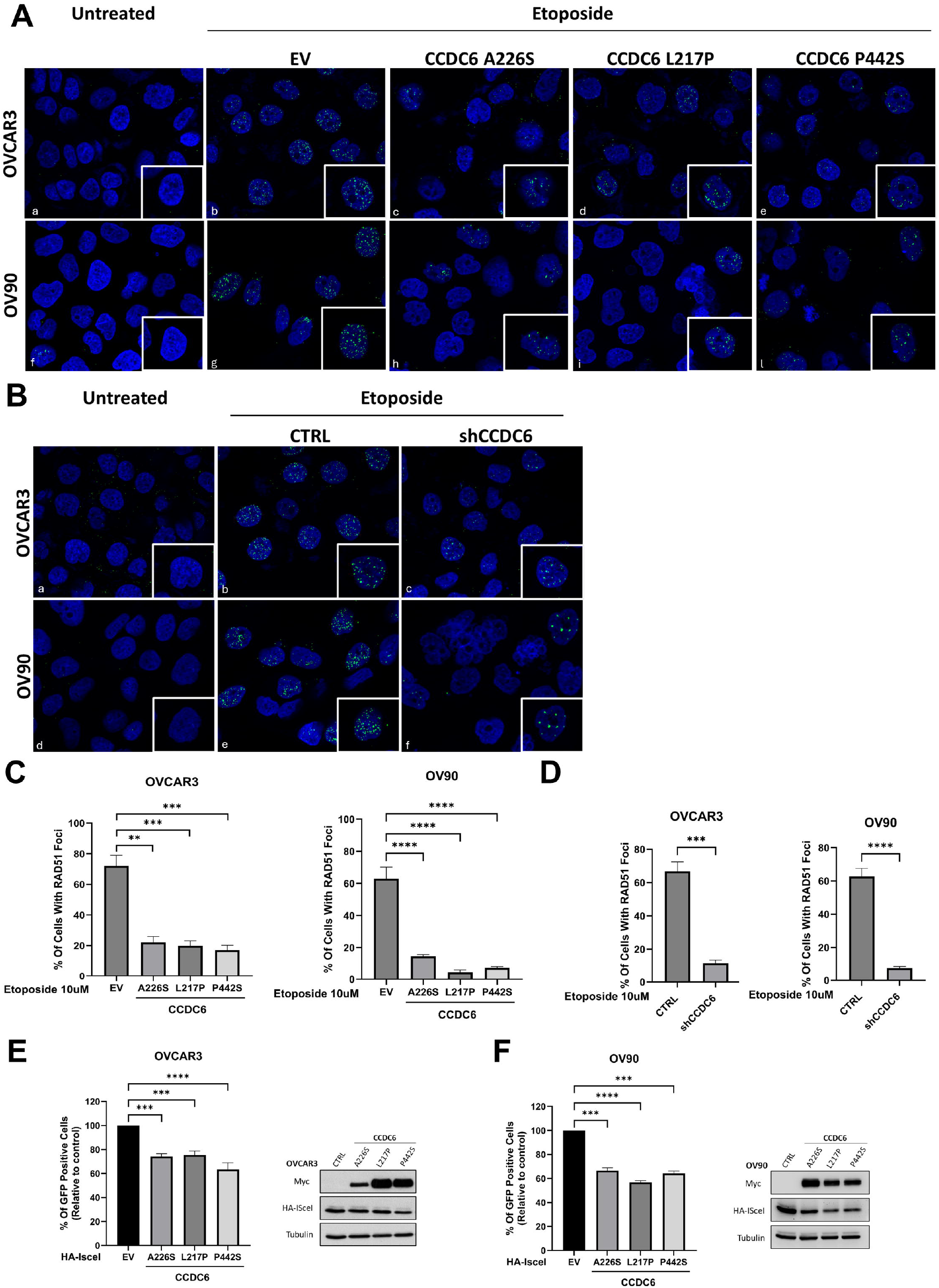
CCDC6 inactivation reduces γH2AX and RAD51 foci formation and reduce the HR repair efficiency in HGSOC cells. (A) Representative immunofluorescence images showing γH2AX nuclear foci formation in OVCAR3 and OV90 cells, following treatment with Olaparib [1μM] for 48 hours. Cells were transfected with the myc-CCDC6 mutated isoforms L217P, A226S and P442, or with the empty vector (EV), as control. (B) Graphs represent the percentage of cells with more than 5 foci. Error bars indicate the standard error mean derived from three independent experiments. Statistical significance was verified by 2-tailed Student’s t-test (^*^ p <0.05; ^**^ p <0.01 and ^***^ p <0.001). (C) RAD51 protein expression was assayed, following cell block procedure, by immunohistochemistry on the HGSOC cell lines OVCAR3 and OV90 stably silenced for CCDC6 (shCCDC6) and control cells (CTRL) (D) RAD51 protein expression was assayed, following cell block procedure, by immunohistochemistry on the HGSOC cell lines OVCAR3 and OV90 expressing Empty Vector (EV) or myc-CCDC6 mutated isoform P442 expression vector (E) RAD51 protein expression levels were detected in OVCAR3 and OV90 cells stably silenced for CCDC6 (shCCDC6) and control cells (CTRL). CCDC6 protein depletion was validated by the anti-CCDC6 antibody. Anti-Tubulin immunoblots are shown as loading control. (F) RAD51 protein expression levels were detected in OVCAR3 and OV90 cells upon transfection with myc-CCDC6 mutated isoforms L217P, A226S and P442, or with the empty vector (EV), as control. Anti-Myc and anti-Tubulin antibodies were employed as transfection and loading controls, respectively (G-H) A) OVCAR3 and OV90 cells were transfected with the mycCCDC6 mutated isoforms (L217P, A226S or P442S) in presence of the DR-GFP construct, and the I-SceI enzyme. The empty vector (EV) was transfected as control. The percentage of GFP positive cells, compared to controls, were plotted as histograms. Error bars indicate the standard error mean derived from three independent experiments. Statistical significance was verified by 2-tailed Student’s t-test (^*^ p <0.05; ^**^ p <0.01 and ^***^ p <0.001). B) Western blot analysis was used to detect the transiently expressed myc-CCDC6 mutated isoforms and the HA-I-SceI. Tubulin is shown as loading control. CCDC6 protein expression was assayed, following cell block procedure, by immunohistochemistry on the ovarian cancer cell lines (a, b) OV90, parental and olaR, (c,d) OVCAR3, parental and olaR, (e, f) PEO1 transiently expressing Empty Vector (EV) or CCDC6 wild type (Myc CCDC6), respectively, (g, h) PEO4 transfected with ShCTRL and ShCCDC6, respectively

To assess if CCDC6 mutants affect HR repair efficiency, we expressed myc-tagged L217P, A226S, and P442S mutants with the DR-GFP reporter and HA-I-SceI endonuclease in OVCAR3 and OV90 cells. After 48 hours, flow cytometry showed fewer fluorescent (recombined) cells in those expressing each CCDC6 mutant compared to control cells (Figure 2E-F). These findings indicate that CCDC6 mutants, similar to CCDC6 depletion, ^21^ impair HR repair in HGSOC cells.

### The CCDC6 mutated isoforms determine PARPi and cisplatin sensitivity in HGSOC cells

HR deficiency predicts PARPi sensitivity and guides treatment decisions. ^4, 29^ We investigated if CCDC6 dysfunction, due to its exclusion from the nucleus, could lead to PARPi sensitivity in HGSOC cells. Expressing CCDC6 mutants in OVCAR3 and OV90 cells increased their sensitivity to Olaparib (Figure 3A-B) and cisplatin (Figure 3C-D). Notably, CCDC6 mutant overexpression further enhanced the synergy between Olaparib and cisplatin (Figure 3E). Our findings suggest that in HGSOC cells, the loss of wild-type CCDC6 nuclear activity in HR creates a synthetic lethal vulnerability to PARP inhibition, contributing to the observed drug synergy.

**Figure 3.**
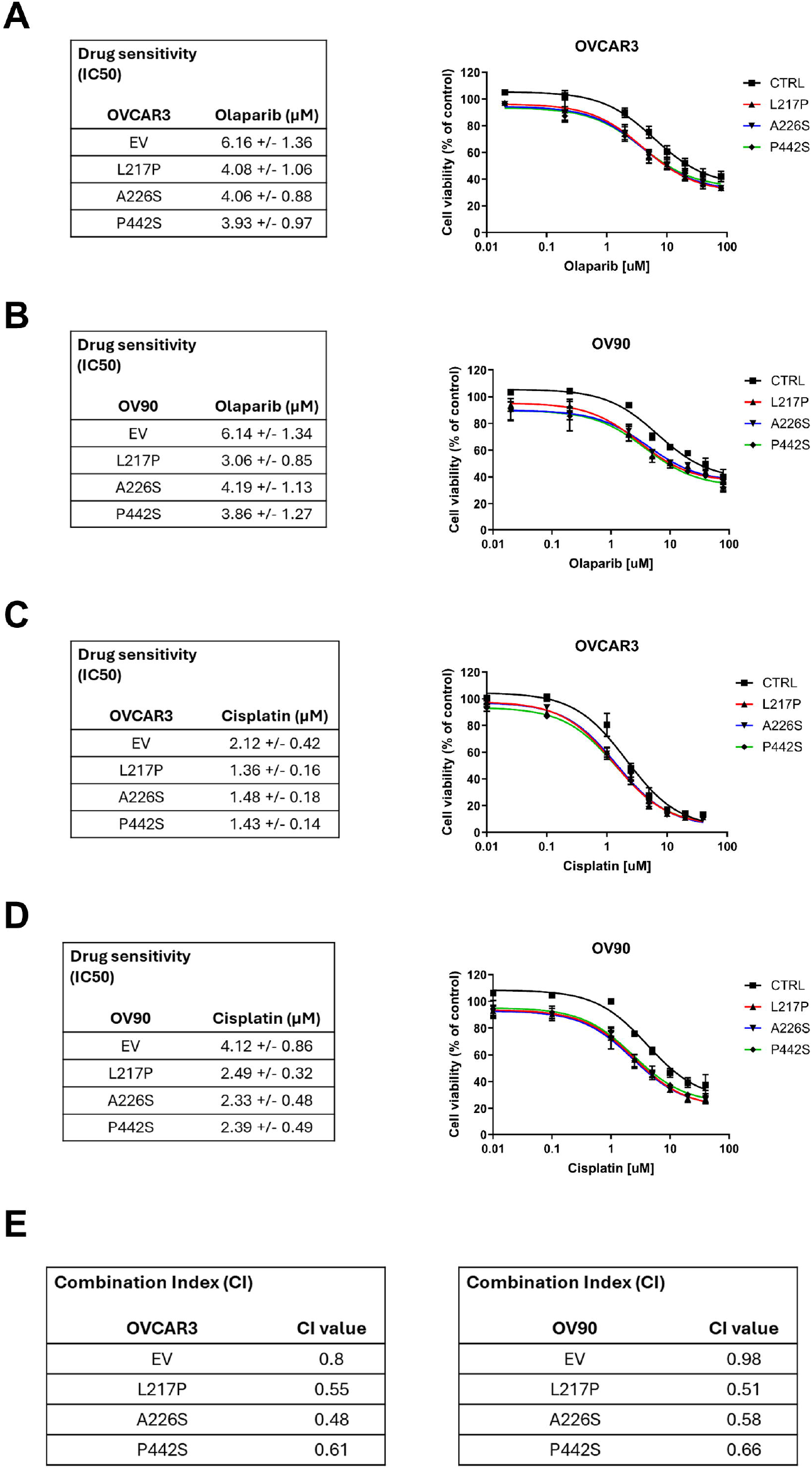
Mutated isoforms of CCDC6 sensitize HGSOC cells to the PARP inhibitor Olaparib alone or in combination with cisplatin. (A, B, C, D) Drugs sensitivity to Olaparib or cisplatin have been evaluated by cell viability assay (Cell Titer 96 Aqueous One Solution assay, Promega) in OVCAR3 and OV90 cell line transfected with the myc-CCDC6 mutated isoforms L217P, A226S and P442 or with the empty vector (EV) exposed to the drug for 144h. The drugs sensitivity is expressed as 50% inhibitory concentration (IC50) values. (E) The combination index values (CI) according to 1:2 concentration ratio of Cisplatin and Olaparib are shown. (CI < 1, CI = 1 and CI > 1 indicate synergism, additive effect and antagonism, respectively).

### The physical or functional loss of CCDC6 confers a HRD phenotype in HGSOC patients from MITO16A

Finally, we explored the link between CCDC6 loss (physical or functional) and HR deficiency (HRD) in HGSOC patients. 185 FFPE tissue samples from the MITO16A trial had been analyzed using genomic and functional (RAD51 foci) HRD tests. ^23, 30^ BRCA1/2 mutational status had also been assessed. ^18, 23^ In these samples we evaluated CCDC6 protein expression and localization via immunohistochemistry and digital image analysis using QuPath. CCDC6 protein levels were quantified using an H-score, which considers both staining intensity and the percentage of positive cells. We then examined the distribution of these H-scores across all samples.

Besides total protein levels, we also analyzed CCDC6’s subcellular localization (nucleus vs. cytosol) to see if its distribution correlates with HRD status. In the MITO16A cohort, 66 of 185 samples (35%) were classified as “CCDC6 inactive” due to either complete nuclear exclusion or very low expression (H-score < 173) (Figure 4A).

**Figure 4.**
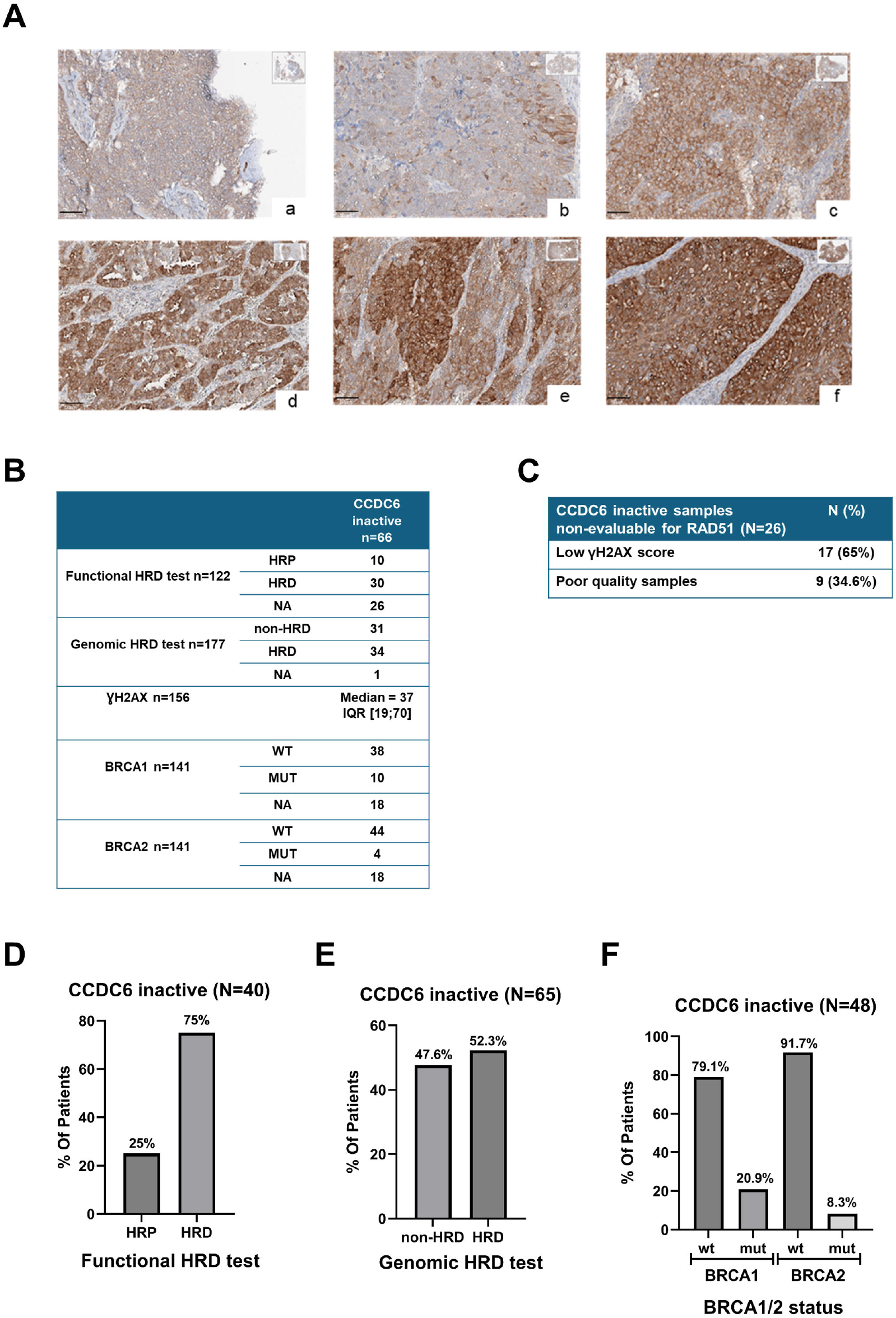
CCDC6 functional inactivation correlate with a HRD phenotype in HGSOC patients. (A) Representative images of immunohistochemical (IHC) analysis of CCDC6 expression (a,b,c) and localization (d,e,f) in FFPE MITO16A patient samples (magnification 40 x) (B) Summary of the MITO16A cases with CCDC6 inactivation that were assessed for HRD test and/or RAD51 test and/or BRCA1/2 status (C) Causes of failure for RAD51 assay in CCDC6 inactive samples (D) Percentages of HRD and non-HRD tumors assessed by genomic or functional HRD testing among CCDC6 inactive MITO16A samples (E) Percentage of BRCA1 wild type or mutated tumors among CCDC inactive MITO16A samples CCDC6 protein expression was assayed, following cell block procedure, by immunohistochemistry on the ovarian cancer cell lines (a, b) OV90, parental and olaR, (c,d) OVCAR3, parental and olaR, (e, f) PEO1 transiently expressing Empty Vector (EV) or CCDC6 wild type (Myc CCDC6), respectively, (g, h) PEO4 transfected with ShCTRL and ShCCDC6, respectively

The association between CCDC6 inactive status and HRD or BRCA mutational status in the MITO16A cohort was summarized in Figure 4B.

The innovative RAD51 immunofluorescence assay was used in the MITO16A trial (185 HGSOC patients) to assess HR repair functionality in patient samples by detecting RAD51 nuclear foci, aiming to correlate it with PARPi sensitivity. ^22, 23^

Interestingly, of the 66 “CCDC6-inactive” samples, 75% (30/40) were HRD-positive by the functional assay (Figure 4D), strongly supporting our in vitro results. Genomic testing showed that 52% (34/65) of CCDC6-inactive samples also had genomic HR deficiency (Figure 4E), with a slightly lower percentage potentially due to test limitations. ^18, 23, 30^

Of note, the “CCDC6 inactive” group distributed mainly in the BRCA1 and BRCA2 wild type patients (Figure 4F), suggesting that CCDC6 impairment may be mutually exclusive with BRCA1/2 mutations.

Our analysis indicates that “CCDC6 inactive” status, reflecting physical or functional CCDC6 loss, is associated with an HRD phenotype. This suggests that PARPi therapy might benefit a wider patient group beyond those with BRCA mutations.

Notably, of the 26 “CCDC6 inactive” samples not analyzed by the RAD51 assay, 17 (65%) showed low γH2AX score (< 25% of geminin-positive cells with γH2AX foci) (Figure 4C), strongly indicating CCDC6 impairment and suggesting these patients also have HRD. Thus, assessing CCDC6 status could identify HRD patients missed by RAD51 assays, potentially broadening PARPi access. Further research is needed to validate this finding.

## Discussion

High-grade serous ovarian carcinoma (HGSOC) patients with BRCA1/2 mutations exhibit homologous recombination deficiency (HRD), leading to sensitivity to PARP inhibitors (PARPi). However, HRD and PARPi sensitivity can occur independently of BRCA mutations.^1-4^ This study identifies loss of CCDC6 function as a novel biomarker and therapeutic target, linking CCDC6 inactivity to HRD and increased PARPi sensitivity. Functional assays and immunohistochemical analysis of MITO16A trial samples revealed that 35% of HGSOC cases exhibited CCDC6 inactivity, with a strong association with HRD by RAD51 and genomic HRD testing. These findings suggest a potential expansion of PARPi eligibility beyond BRCA-mutant patients.

CCDC6 has been proposed as a tumor suppressor gene due to its role in the surveillance of DNA integrity.^20, 26^ Preclinical data demonstrated an association between CCDC6 gene product depletion in HGSOC cells and Olaparib sensitivity suggesting CCDC6 as a potential biomarker of BRCAness phenotype and PARPi sensitivity.^21^

The easier accessibility of NGS technology and advancements in diagnostic tools have led to a significant increase in omics data, including extensive information on CCDC6 molecular alterations in online databases. These alterations are highly heterogeneous, involving fusions with various partners and point mutations clustered in different coding regions. A systematic characterization of CCDC6 alterations in HGSOC was lacking, and we questioned if these could cause a CCDC6 loss-of-function phenotype.

The characterized CCDC6 missense mutations, found heterozygously in HGSOC, produce a phenotypic effect similar to that of native protein depletion. Furthermore, our data elucidate the mechanism, as overexpressed mutant CCDC6 isoforms cause nuclear exclusion of the wild-type protein, leading to a CCDC6 loss of function not explained by the Knudson two-hit theory. This dominant-negative behavior has been previously observed with truncated and mutated CCDC6 isoforms in other cancer cell models. ^25^

The CCDC6 mutants L217P, A226S, and P442S retain the ability to heterodimerize with wild-type CCDC6, causing altered subcellular localization and impaired native protein function. Immunofluorescence showed reduced nuclear localization of wild-type CCDC6 with these mutants, indicating that cytosolic delocalization/nuclear exclusion disrupts the CCDC6-PP4c interaction. ^20, 21^ This disruption leads to unrestrained PP4c activity, affecting H2AX phosphorylation, DNA damage signaling, cell cycle progression, and HR-mediated DSB repair, as evidenced by a decrease in GFP-positive cells in the HR DR-GFP reporter assay upon mutant expression.

Given that CCDC6 mutants primarily delocalize the native protein to the cytosol, we emphasized the importance of evaluating CCDC6 intracellular localization (cytosol/nucleus), in conjunction with CCDC6 protein intensity, in HGSOC patient tissue samples. This assessment was performed via immunohistochemistry on a large cohort of HGSOC specimens from the MITO16A clinical study.^23^ We classified 66 out of 185 analyzed samples (35%) as ‘CCDC6 inactive’, based on either complete nuclear exclusion or an H-score within the first quartile (H-score < 173), signifying null or minimal CCDC6 expression (Figure S1B).

Consistent with our in vitro observation that CCDC6 loss or nuclear exclusion reduces RAD51 and γH2AX positive cells and impaired HR repair, MITO16A patients with ‘CCDC6 inactive’ tumors predominantly showed HR impairment, as confirmed by genomic HRD testing and, notably, the RAD51 functional assay.

However, the RAD51 assay had a 30% failure rate due to sample issues or low DNA damage (ie low γH2AX score).^23^ Considering that CCDC6 physical or functional impairment significantly reduces γH2AX foci formation, we conclude that ‘CCDC6 inactive’ status appears to be a valuable tool to identify HRD patients eligible for PARPi therapy who might otherwise be excluded based solely on RAD51 assay unsuitability. Additional investigation is necessary to reinforce this observation.

Overall, the CCDC6 mutated isoforms, behaving in a dominant negative manner versus the CCDC6 native protein, confer to the tumoral cells an increased genomic instability, due to the defective HR, and a vulnerability exploitable by precision medicine approaches, as shown in vitro by enhanced sensitivity to PARPi treatment and verified by cell viability assays. The enhanced cytotoxic effect of Olaparib and the increased synergistic effect of PARPi when combined with other DNA damaging agents, such as Cisplatin, is in support of a BRCAness phenotype dependent on CCDC6 mutated isoforms, that we have characterized.

Here, we’ve learned that DNA mutations can mislocalize CCDC6, impairing its nuclear function in HGSOC. Future work should explore other mechanisms, including CCDC6 interaction with nuclear import/export proteins like XPO1.^31^ Changes in CCDC6 post-translational modification sites (by kinases, phosphatases, or stability enzymes) might also increase turnover, alter dynamics, and affect protein complex interactions.^20, 26^ Furthermore, CCDC6’s role in DDR and HR isn’t fully understood. Computational analysis suggests CCDC6 interacts with HR proteins like BRCC3 and BAP1 (of the BRCA1 complex) ^32^, and ongoing work explores a potential interaction with PALB2. Future studies will clarify the biochemical dynamics of CCDC6 interactions (native and mislocalized forms) under various conditions. Beyond point mutations, uncharacterized gene rearrangements, such as the CCDC6-ANK3 fusion in HGSOC ^33^, also warrant investigation.

Notably, the observation that the CCDC6 mutants have the ability to mainly delocalize the native protein in the cytosol suggested, beside the intensity, a careful evaluation of the CCDC6 intracellular localization (cytosol/nucleus) in tissue samples of HGSOC patients. In this work, this kind of evaluation has been pursued by immunohistochemistry in a large series of HGSOC specimens obtained by the MITO16A clinical trial that also provided patients information about genomic and functional HRD tests.

It is of interest that the study demonstrated a significant association between CCDC6 inactivity and HRD, as evidenced by both genomic and functional analyses. Of particular significance is the observation that this association was maintained in the presence of wild-type BRCA1/BRCA2 genes, suggesting a crucial role for CCDC6 in DNA repair pathways that are independent of these well-characterized genes.

However, the MITO16A study primarily aimed to identify clinical and biological factors predicting progression-free or overall survival in patients receiving first-line chemotherapy and bevacizumab, rather than focusing on PARPi response. We are currently investigating CCDC6 behavior in BRCA wild-type patients with newly diagnosed advanced epithelial ovarian cancer who achieved a partial or complete response to first-line platinum-based chemotherapy and are enrolled in the multicenter, prospective, single-arm MITO35A trial of Olaparib maintenance therapy. Thus, the MITO35a findings could translate MITO16A’s initial observations into clinical practice.

In conclusion, our findings establish CCDC6 inactivity, defined by reduced expression or nuclear exclusion, as a novel and clinically actionable biomarker for homologous recombination deficiency (HRD) in high-grade serous ovarian carcinoma (HGSOC). Importantly, CCDC6 status identifies HRD-positive tumors even in the presence of wild-type BRCA1/2 genes, thereby expanding the population of patients potentially eligible for PARPi therapy. The combination of CCDC6 immunohistochemical assessment with functional RAD51 assays provides a promising strategy to refine patient selection for precision medicine approaches. Future validation in independent cohorts and prospective clinical trials will be essential to integrate CCDC6 evaluation into standard HRD diagnostic workflows, with the ultimate goal of improving therapeutic outcomes for a broader subset of HGSOC patients.

## Aknowledgements

Our investigations have been financed by “Progetto NUTRAGE”, CNR to AC, CNR – IEOMI, “Progetto POR FESR 2014–2020 TECHNOLOGY PLATFORM” LOTTA ALLE PATOLOGIE ONCOLOGICHE Sviluppo di Approcci Terapeutici INnovativi per patologie neoplastiche resistenti ai trattamenti (SATIN), to AC, CNR – IEOMI, and “Progetto SerGenCovid”, (CNR, FOE 2021) to AC, CNR – IEOMI. We acknowledge financial support under the National Recovery and Resilience Plan (NRRP), Mission 4, Component 2, Investment 1.1, Call for tender No. 1409 published on 14.9.2022 by the Italian Ministry of University and Research (MUR), funded by the European Union – NextGenerationEU – Prot. P2022TPPZL, CUP B53D2303323 0001, Grant Assignment Decree No. 1317, adopted on 08/08/2023 by the Italian Ministry of Ministry of University and Research (MUR) to AC, CNR – IEOMI. The study was also partially supported by AIRC IG18921 and IG25932 to SP, Ministry of Health RF-2016-02363995 to SP, Ministry of Health PNRR-MAD-2022-12375663 to SP, and Ministry of Health Ricerca Corrente L4/81_25 to SP.

We are grateful to the Flow Cytometry facility, funded by “Progetto Dipartimento di Eccellenza 2018–2022, Legge 11 dicembre 2016, n. 232” to the Department of Molecular Medicine and Medical Biotechnology, University of Naples Federico II, Naples, Italy, for supporting flow cytometry experiments for the current study.

The authors also thank the cell culture facility of CEINGE-Advanced Biotechnologies “Franco Salvatore”, Naples, Italy.

## Declaration of Generative AI and AI-assisted technologies in the writing process

Google Gemini 1.5 Pro was used to help to revise the text written by the authors, in order to enhance the grammar and English language of this work. The authors then critically reviewed and revised the output, ensuring full responsibility for the content.

## Author contributions

DC: Conceptualization, Data curation, Formal analysis, Investigation, Supervision, Writing – review & editing, FM: Data curation, Formal analysis, Methodology, Validation, Visualization, Writing – review & editing, BP: Data curation, Methodology, Validation, Visualization, LR: Formal analysis, Investigation, Methodology, SV: Investigation, Methodology, Visualization, SCC: Investigation, Methodology, Visualization, IDB: Formal analysis, Data curation, Software, Validation, DC: Data curation, Formal analysis, Resources, Validation, Visualization, RC: Investigation, Methodology, Project administration, CB: Investigation, Methodology, CN: Data curation, Formal analysis, Validation, EP: Data curation, Formal analysis, DK: Resources, EDC: Data curation, Formal analysis, Methodology, Validation, GLS: Data curation, Software, Methodology, SM: Data curation, Formal analysis, Methodology, Software, DR: Investigation, Methodology, AS: Investigation, Methodology, Visualization, LA: Data curation, Formal analysis, Methodology, Software, FM: Data curation, Investigation, Methodology, Validation, MM: Investigation, MAV: Methodology, Validation, AI: Formal analysis, Methodology, Visualization, FP: Data curation, Formal analysis, Supervision, SP: Conceptualization, Formal analysis, Resources, Supervision, Writing – review & editing, AC: Conceptualization, Data curation, Formal analysis, Supervision, Writing – original draft, Writing – review & editing

## Abbreviations

HGSOC: High Grade Serous Ovarian Cancer
DSB: Double Strand Break
HR: Homologous Recombination
PARPi: poly-(ADP-ribose)-polymerase inhibitor
PFS: Progression-free survival
HRD: Homologous Recombination Deficiency
CCDC6: Coiled Coil Domain Containing 6
FFPE: Formalin Fixed Paraffin Embedded
MITO: Multicenter Italian Trials in Ovarian cancer
OC: Ovarian carcinoma
NSCLC: Non-Small Cell Lung Cancer

---

**Figure SSI:**
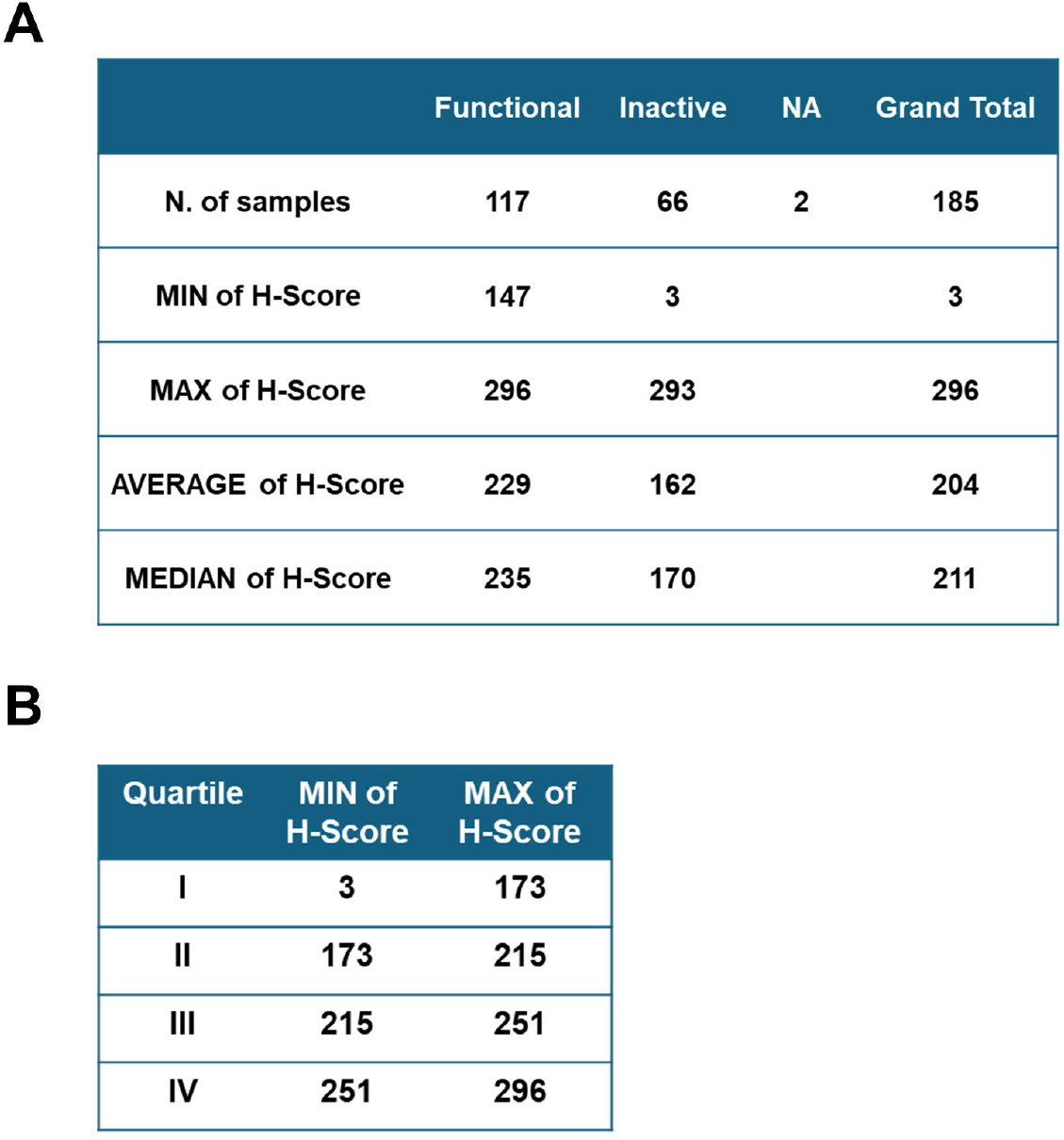
(A) Summary statistics of H-scores for protein localization categories. Functional. Inactive, and Not Analysed (NA) samples arc presented, including the number of samples (N), minimum, maximum, average, and median H-scores. (B) Distribution of IHC H-scores for CCDC6 within the M1T016A cohort.

**Figure S2:**
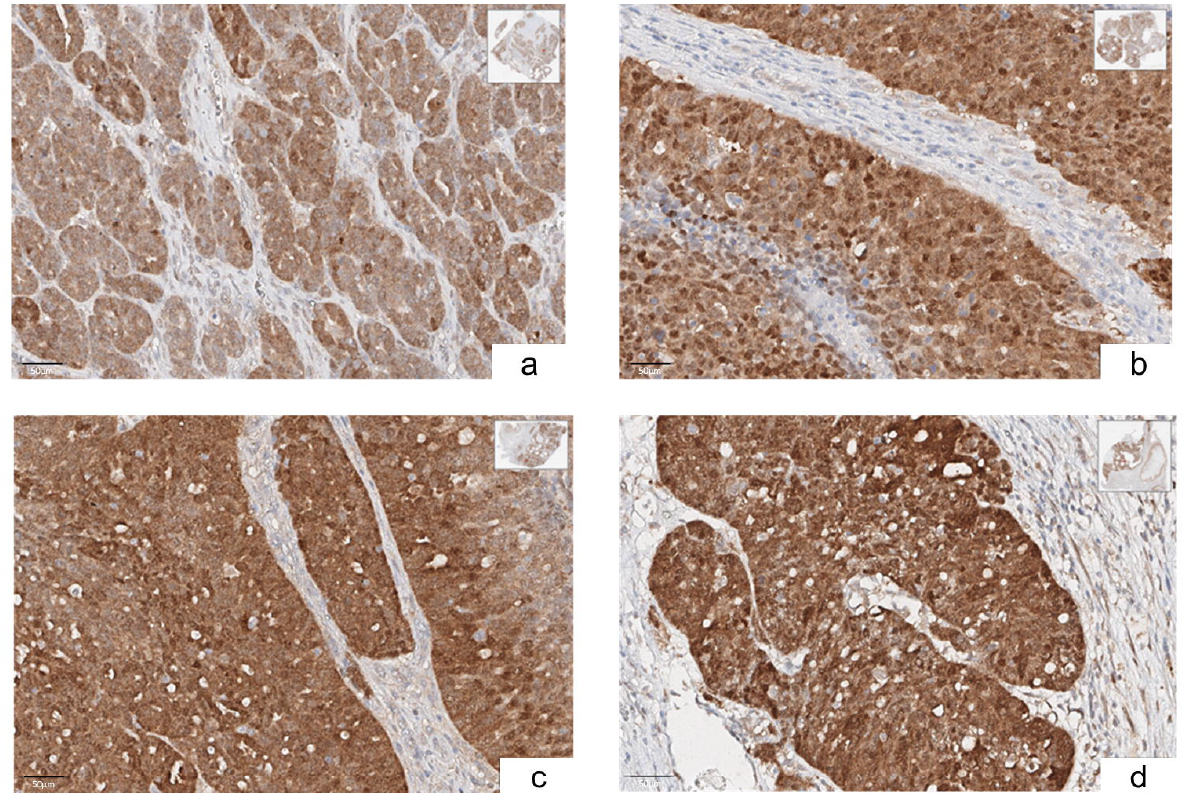
Immunohistochemical (IHC) analysis of CCDC6 staining intensity (H-score >173) and localization (diffuse or mainly nuclear), identified the “CCDC6 functional” MITO 16A patient set (N = 117) (Magnification 40X).

**Figure S3:**
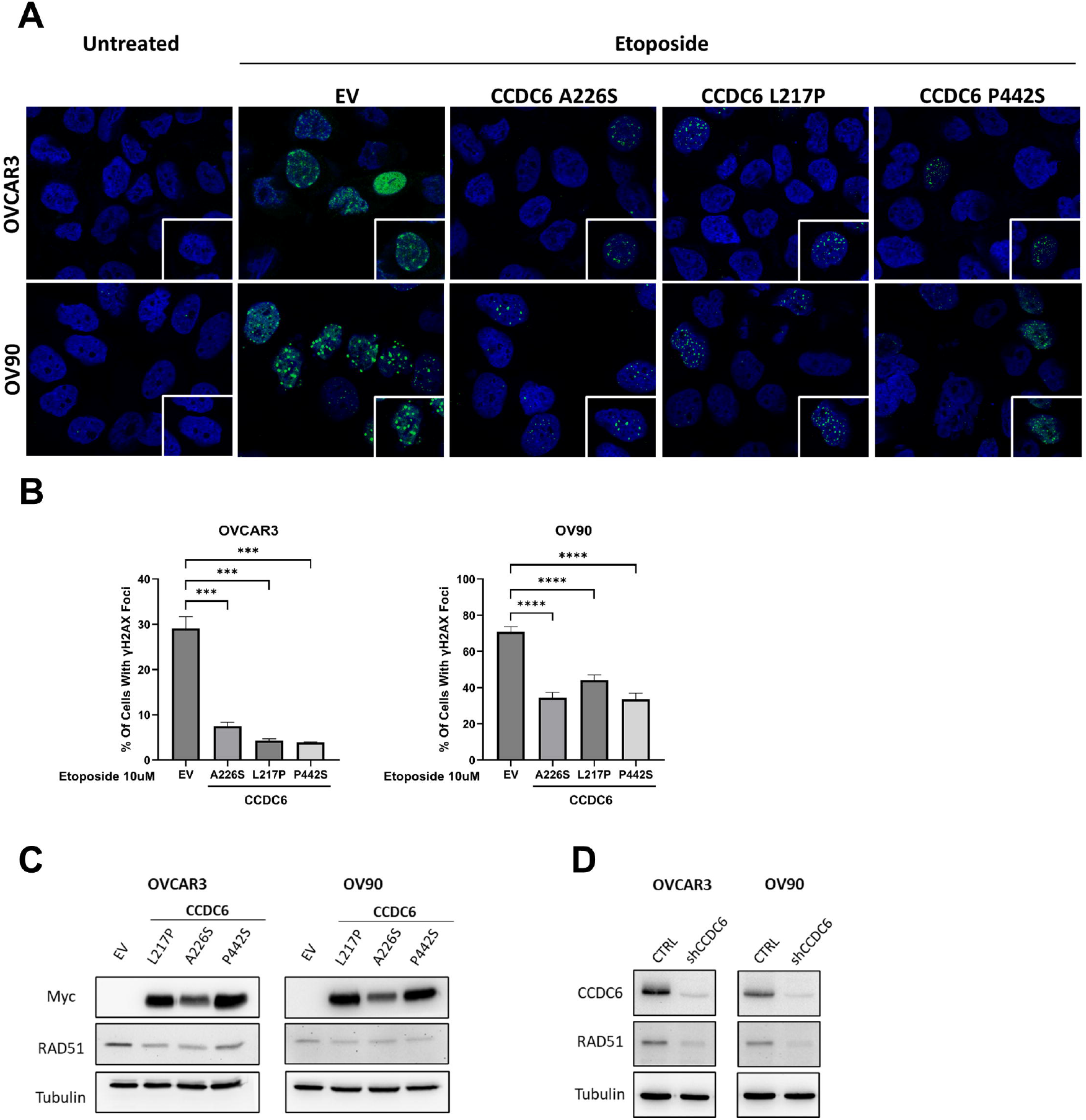
(A) Representative immunofluorescence images showing γH2AX nuclear foci formation in OVCAR3 and OV90 cells, following treatment with Etoposide [10 μM] for 8 hours. Cells were transfected with the myc-CCDC6 mutated isoforms L217P. A226S and P442S, or with the empty vector (EV), as control. (B) Graphs represent the percentage of cells with more than 5 foci. Error bars indicate the standard error mean derived from three independent experiments. Statistical significance was verified by 2-tailcd Student’s t-test (* p <0.05; ** p <0.01; *** p <0.001 and **** p <0.0001). (C) RAD51 protein expression levels were detected in OVCAR3 and OV90 cells upon transfection with myc-CCDC6 mutated isoforms L217P, A226S and P442S, or with the empty vector (EV), as control. Anti-Myc and anti-Tubulin antibodies were employed as transfection and loading controls, respectively. (D) RAD51 protein expression levels were detected in OVCAR3 and OV90 cells stably silenced for CCDC6 (shCCDC6) and control cells (CTRL). CCDC6 protein depletion was validated by the anti-CCDC6 antibody. Anti-Tubulin immunoblots are shown as loading control.

